# Commonly used organoid culture media prevent ferroptosis

**DOI:** 10.1101/2024.04.29.591759

**Authors:** John K. Eaton, Priya Chatterji, Yan Y. Sweat, Rachelle A. Victorio, Mathias J. Wawer, Vasanthi S. Viswanathan

## Abstract

Organoids enable the recreation of organ physiology *in vitro* and serve as powerful models for biology in basic life science research and drug discovery and development. However, organoid culture requires complex media formulations that include antioxidants, potentially confounding experimental results sensitive to such conditions. Here we report that the growth conditions used commonly to generate organoid models inhibit ferroptosis, an iron-dependent form of lipid peroxidative cell death with relevance to human disease, thus rendering such models incompatible with ferroptosis research. We identify medium components that diminish or eliminate ferroptosis sensitivity and outline strategies for avoiding anti-ferroptotic culture conditions in organoid and other cell culture applications. These findings provide a roadmap for adapting organoid models for the study of ferroptosis and leveraging their strengths for advancing ferroptosis-modulating therapeutics.

## Introduction

Ferroptosis is a disease-relevant form of cell death characterized by the iron-dependent formation of toxic lipid peroxides and subsequent loss of membrane integrity.^1–3^ Modulation of ferroptosis holds immense potential for impacting human health across several therapeutic areas if translated successfully in the pre-clinical setting. The adoption of organoid systems, which constitute an important translational bridge between two-dimensional (2D) *in* vitro cell culture systems and three-dimensional (3D) *in vivo* contexts will be an important and necessary aspect of building pre-clinical and early-development workflows for ferroptosis translation. Such innovation will, however, pose certain challenges. Like many biological processes, the capacity of cells to undergo ferroptotic cell death can be influenced by environmental conditions, including the availability of substrates required for lipid peroxidation (polyunsaturated lipids, oxygen and iron)^1,4–6^ and factors influencing cellular antioxidant defenses.^7–15^ To correctly perform and interpret ferroptosis-related experiments requires knowledge of and control over experimental parameters impacting ferroptosis and their physiological relevance.

Medium formulations used to generate organoids are more complex than media typically used in 2D cell culture and often contain dozens of additional components including proteins and small molecules.^16^ In fact, the development of such complex culture media formulations was critical to the success of organoid technology through the inclusion of growth factors and other nutrients that enable 3D cell growth *in vitro*.^17,18^ Many organoid and tumoroid medium formulations that enable model generation from specific cells or tissue types have been described in detail.^16,19^ However, media formulations offered by commercial vendors are often proprietary and do not disclose the identity or concentration of components. How any of these known and unknown ingredients may confound ferroptosis measurements and impede the use of organoids for ferroptosis pre-clinical research is not established.

Here we report that frequently used organoid media contain a variety of components that can suppress ferroptosis. The components include but are not limited to the ferroptosis-inhibiting antioxidant vitamin E.^20^ Our findings suggest that caution must be exercised when probing ferroptosis in systems using undefined media formulations like those often employed in the study of organoid models. We additionally provide recommendations for characterizing cell culture conditions to determine suitability for studying ferroptosis.

### Commercial organoid media inhibit ferroptosis in 2D and 3D cell culture formats

We tested a commercial organoid medium formulation to determine its effect on ferroptosis sensitivity of 2D and 3D cell culture models. The complete identity and concentration of components in this medium have not been disclosed by the manufacturers and, to our knowledge, the impact of these culture conditions on cellular ferroptosis sensitivity has not been investigated systematically. We initially selected cell lines sensitive to ferroptosis in 2D formats and cultured these cells with either standard culture media or commercial organoid media. Cells were then treated with a panel of small molecules including ferroptosis inducers of distinct structural and mechanistic classes, non-ferroptosis inducing molecules known to generate reactive oxygen species (ROS), and other cytotoxic agents (Supplementary Figure 1). In multiple cell lines, we observed that organoid media completely suppressed the effects of ferroptosis inducers but had no effect on lethal compounds that do not promote ferroptosis (Figures 1A, 1B, and 1C; Supplementary Figures 2–4). Organoid medium was as effective at protecting cells from ferroptosis as the well-validated ferroptosis inhibitor ferrostatin-1 (fer-1), a small molecule with radical-trapping antioxidant activity.^1,21,22^

**Figure 1:**
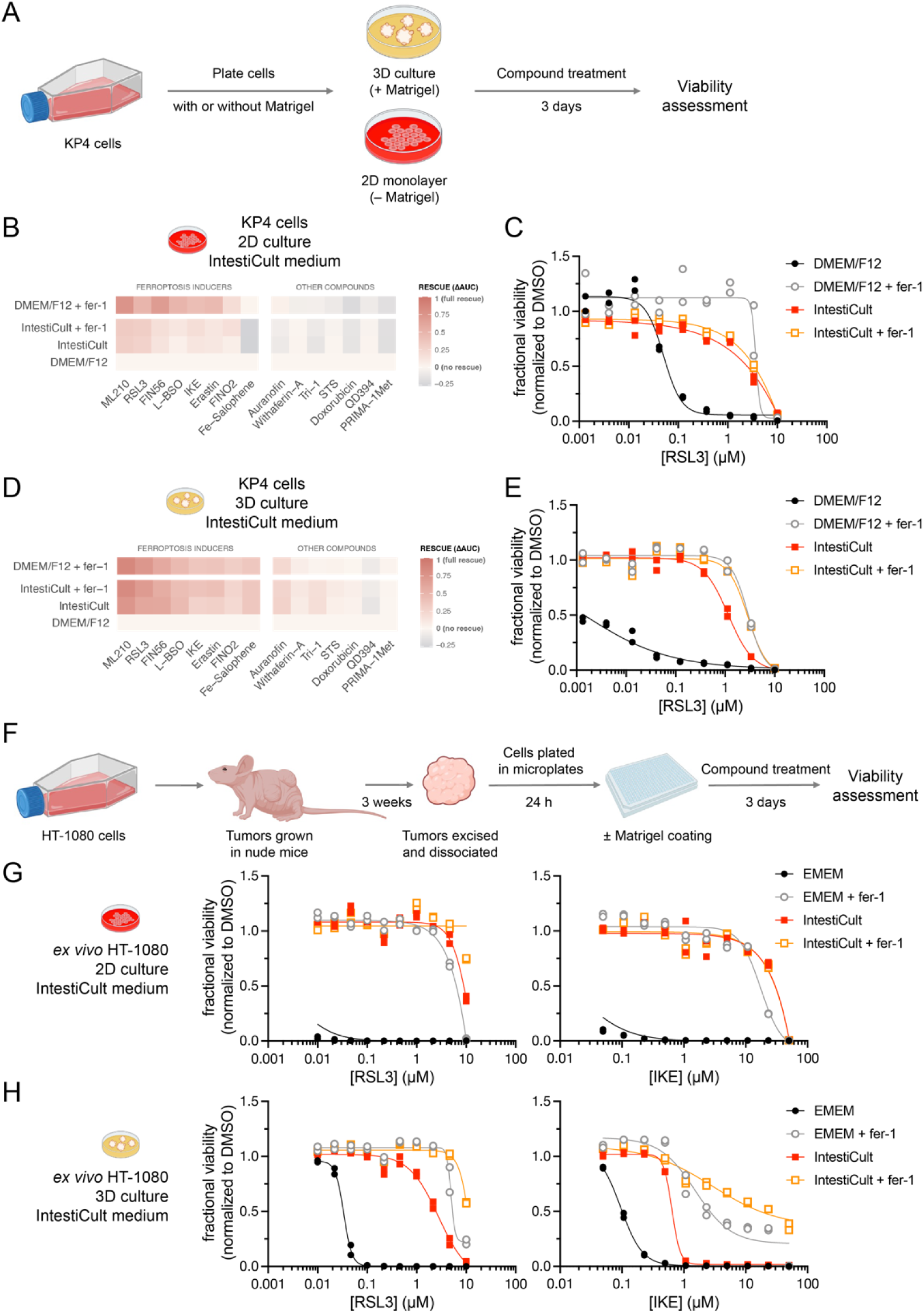
Commercial organoid medium suppresses ferroptosis. (A) Scheme depicting the assessment of organoid media on the ferroptosis sensitivity of cancer cell lines in 2D and 3D formats. (B) IntestiCult™ medium protects KP4 cells from ferroptosis inducers in 2D culture. Heatmap colors are based on the difference in area-under-the-curve (ΔAUC) metric for viability of compound-treated cells relative to the DMEM/F12 medium condition. AUC values for each compound relative to vehicle-treatment are derived from a 12-point dose response curve, n = 2 technical replicates. Compound treatment was performed for 72 h. (C) Dose response curves for RSL3 treatment from panel B. Data are plotted as two individual technical replicates. See also Supplementary Figure 2A. (D) IntestiCult™ medium protects KP4 cells from ferroptosis inducers in 3D culture. Heatmap is plotted as described for panel B. (E) Dose response curves for RSL3 treatment from panel D. Data are plotted as two individual technical replicates. See also Supplementary Figure 2B. (F) Scheme depicting the *ex vivo* assessment of HT-1080 cells in 2D and 3D culture formats. (G and H) IntestiCult™ medium prevents ferroptosis in *ex vivo* HT-1080 cells plated in 2D (panel G) and 3D (panel H) culture formats. Cells were treated with the indicated compounds for 72 h. Data are plotted as two individual technical replicates.

Previous reports have suggested that ferroptosis vulnerability may be impeded by cellular contacts, such as those modeled in 3D culture,^23^ so we next investigated whether the effects on ferroptosis sensitivity in 3D growth format might be a function of media conditions rather than 3D format per se. Our results demonstrate that ferroptosis sensitivity was preserved in all tested 3D formats when using standard 2D culture medium (Figures 1A, 1D, and 1E; Supplementary Figures 2 and 4). By contrast, cells were impervious to all ferroptosis inducers when cultured in 3D format with commercial organoid media, just as we observed in the 2D format (Figures 1D and 1E; Supplementary Figures 2–4). Importantly, the Matrigel matrix used to grow cells in 3D did not impede sensitivity to ferroptosis (Figure 1 and Supplementary Figure 4). However, it is possible that other matrices used for 3D culture may contain ferroptosis-modulating components. Commercial organoid media also prevented ferroptosis *ex vivo* in cells derived from human xenografted tumors when cultured in both 2D and 3D formats (Figures 1F, 1G, and 1H). These same xenograft-derived tumor cells underwent robust ferroptosis *ex vivo* when cultured in normal basal media. Taken together, these observations demonstrate that components of organoid media potently inhibit ferroptosis and it is therefore important to perform appropriate validation when using models with these media for studying ferroptosis.

### Common organoid medium supplements include ferroptosis-inhibiting components

A variety of culture conditions have been reported for organoids depending on cell origin and desired model properties.^16^ Given the complexity of such media and the lack of verifiable composition information for many commercial options, we focused on surveying examples from three classes of common additives included in literature medium recipes: commercial supplement mixtures (e.g., B27), small-molecule pathway inhibitors (e.g., ROCK inhibitors), and nutrients (e.g., N-acetylcysteine). We tested these additives by adding them individually to 2D culture medium in which cells retain ferroptosis sensitivity and measuring their effect on cell viability across a panel of structurally and mechanistically diverse ferroptosis inducers and other lethal small molecules.

### B27 supplement contains vitamin E at a concentration sufficient to prevent ferroptosis

We initially investigated supplement mixtures B27 and N-2, which are widely included in organoid medium formulations and contain multiple components that may potentially influence cellular ferroptosis sensitivity (Supplementary Tables 1 and 2). N-2 supplement was developed for the serum-free culture of neuroblastoma cells and contains five components^24^ at concentrations specified by commercial vendors (Supplementary Table 2). B27 is a mixture of 20 proteins and small molecules (Supplementary Table 1) originally developed to culture hippocampal neurons.^25,26^ B27 is known to contain multiple antioxidants, including the ferroptosis inhibitor vitamin E, but vendors do not disclose the specific concentrations of these components.^27^ Inclusion of B27 supplement in 2D cell culture medium conferred protection to multiple cell lines against ferroptosis-inducing molecules but not to other lethal compounds (Figures 2A and 2B; Supplementary Figure 5). Medium supplemented with B27 was also sufficient to prevent the formation of lipid hydroperoxides in cells as assessed using a C11-BODIPY peroxidation sensor (Figure 2C). As expected, based on the above ferroptosis-specific effects of B27, supplementation with B27 did not affect the compound sensitivity of a ferroptosis-insensitive cell line (Supplementary Figure 6).

**Figure 2:**
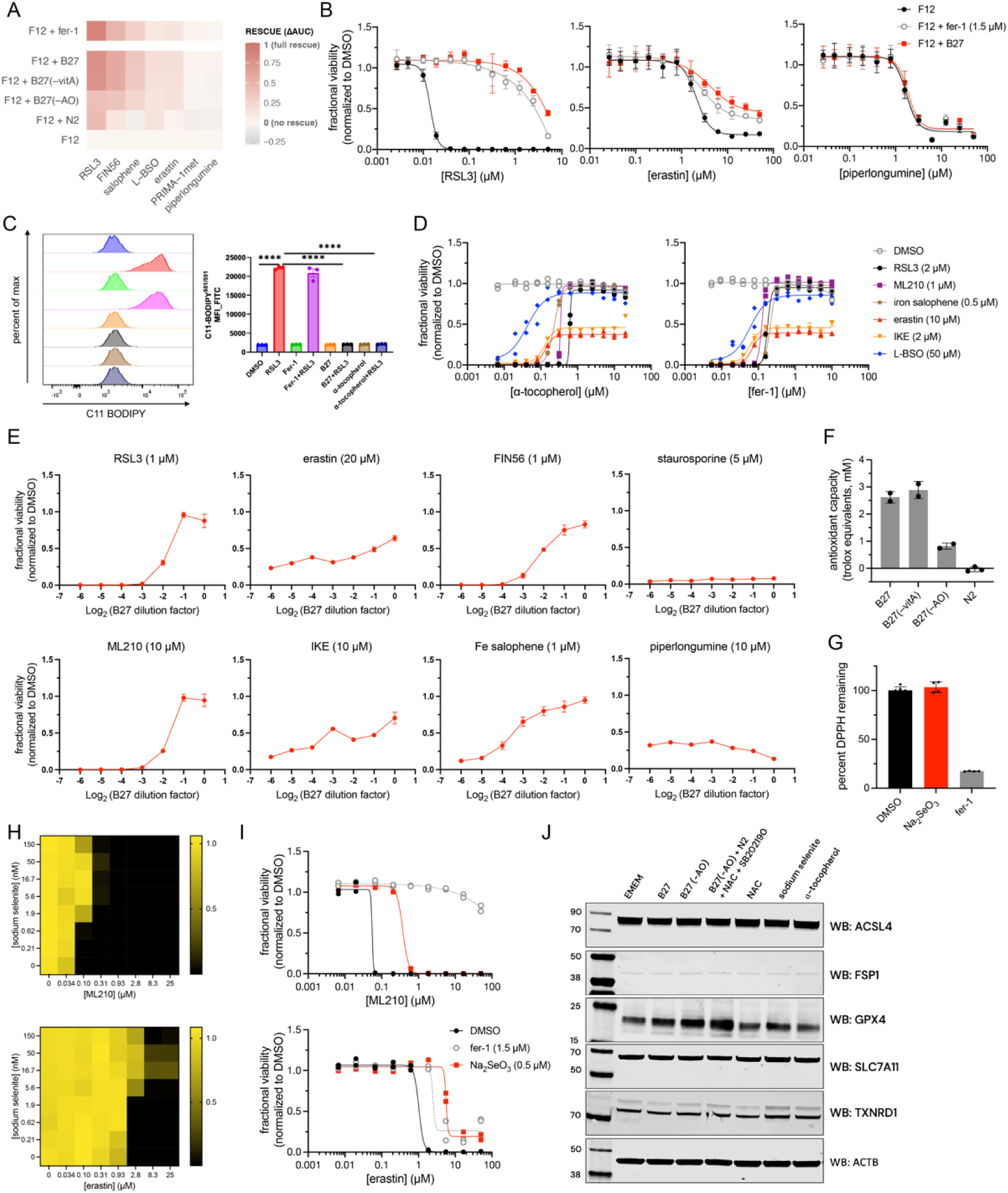
B27 and N-2, common organoid medium supplements, suppress ferroptosis. (A) RKN cells cultured in F12 medium supplemented with B27 or N-2 are protected from ferroptosis inducers. Heatmap colors are based on the difference in area-under-the-curve (ΔAUC) metric for viability of compound-treated cells relative to the F12 medium condition. AUC values for each compound relative to vehicle-treatment are derived from 12-point dose response curves with n = 3 technical replicates. Compound treatment was performed for 72 h. (B) Selected dose response curves from the heatmap in panel A. Data are plotted as mean ± s.d. for n = 3 technical replicates. See also Supplementary Figure 7. (C) B27 supplement suppresses lipid peroxidation in HT-1080 cells as assessed by C11-BODIPY 581/591. Data are plotted as mean ± s.d. for n = 3 replicates. (D) Dose-response assessment of ferroptosis suppression by ɑ-tocopherol and ferrostatin-1 in RKN cells. Cells were treated for 24 h with the indicated ferroptosis inhibitor and then incubated with ferroptosis inducers for 72 h prior to viability assessment. Data are plotted as mean ± s.d. for n = 3 technical replicates. (E) Dilution-dependent assessment of ferroptosis suppression by B27 supplement. RKN cells were treated with the indicated compounds for 72 h with varying dilutions of B27. Data are plotted as mean ± s.d. for n = 6 technical replicates. (F) N-2 supplement does not exhibit direct antioxidant activity. Data are plotted as n ≥ 2 individual technical replicates. (G) Sodium selenite does not exhibit radical trapping activity in the DPPH assay. Data are plotted as mean ± s.d. for n ≥ 4 technical replicates. (H) Sodium selenite protects RKN cells in a dose-dependent manner from ferroptosis inducers. Each well represents the fractional cell viability of cells subjected to the indicated condition relative to DMSO treatment. (I) Dose-response plots demonstrating rescue of ferroptosis inducers by sodium selenite. Cells were incubated with sodium selenite for 24 h before treatment with test compounds for 72 h. Data are plotted as two individual technical replicates. (J) Medium supplements containing sodium selenite increase GPX4 protein levels in RKN cells but do not alter the expression of other ferroptosis-relevant proteins as assessed by western blot. Supplements were used at 1x dilution in F12 medium. Small molecules were added at indicated concentrations: sodium selenite (0.1 µM), SB 202190 (10 µM), ɑ-tocopherol (2 µM), N-acetylcysteine (NAC, 1.25 mM).

At the recommended dilution of B27 in published organoid medium formulations, the final concentration of vitamin E is over 4 µM in the form of ɑ-tocopherol and the prodrug analog ɑ-tocopherol acetate (Supplementary Table 1). At this concentration, ɑ-tocopherol was sufficient to completely protect multiple cell lines from small-molecule ferroptosis inducers to an extent comparable with other ferroptosis inhibitors, including ferrostatin-1 (Figure 2D). This concentration far exceeds the level of vitamin E from fetal bovine serum (FBS) commonly added to cell culture media, which is approximately 45 nM when used at a typical FBS concentration of 10% by volume.^28^ Consistent with the potency of ɑ-tocopherol for ferroptosis rescue, B27 supplement can be diluted several fold and still prevent ferroptosis inducers from killing cells (Figure 2E). The ability of ɑ-tocopherol to rescue cells from ferroptosis varies slightly across different classes of ferroptosis inducers. For example, ferroptosis inducers that inhibit glutathione biosynthesis, a ferroptosis induction target we recently validated,^29^ were rescued more potently by ɑ-tocopherol than by other radical-trapping antioxidants or iron chelators (Figure 2D).

### Selenium present in B27 and N-2 supplements suppresses ferroptosis sensitivity

We next sought to test whether B27 contains additional components beyond vitamin E that inhibit ferroptosis. Compared to standard B27, commercially available “antioxidant-free” B27 (B27–AO) lacks ɑ-tocopherol, ɑ-tocopherol acetate, superoxide dismutase, catalase, and glutathione (Supplementary Table 1). However, B27–AO was still able to partially protect cells from ferroptosis inducers (Figure 2A and Supplementary Figure 7). Likewise, N-2 supplement lacks vitamin E (Table 2) yet protects cells against certain ferroptosis inducers to an extent comparable to B27– AO (Figure 2A and Supplementary Figure 7).

To isolate components in B27-AO and N-2 supplements that suppress ferroptosis, we focused our attention on the five shared ingredients between N-2 and B27–AO supplements (Supplementary Tables 1 and 2). Among these, sodium selenite (Na_2_SeO_3_) is a major source of selenium for cells under standard *in vitro* culture conditions. Selenite, like N-2 supplement, does not possess direct antioxidant activity (Figures 2F and 2G). However, it has been reported to protect cells from ferroptosis-inducing perturbations both through increasing GPX4 expression^7,30–32^ and by generating hydrogen selenide (H_2_Se), which acts as an antioxidant in a selenoprotein-independent manner.^33^ Indeed, we found that addition of sodium selenite to cell culture media partially protected cells from all ferroptosis inducers tested and did so at concentrations comparable to those found in B27 and N-2 supplements (Figures 2HC and 2ID). Addition of sodium selenite did not affect the potency of other lethal compounds (Supplementary Figure 8). We further observed that cells cultured in medium supplemented with either B27–AO, N-2, or sodium selenite had elevated levels of GPX4 protein when compared to cells cultured in basal medium (Figure 2J). The expression of other ferroptosis-relevant proteins was unaffected (Figure 2J). These findings demonstrate that the levels of selenium present in B27 and N-2 supplements alone are sufficient to impact ferroptosis sensitivity, at least in part through GPX4 upregulation. When combined with other media components that act to inhibit ferroptosis through other distinct mechanisms, further synergistic anti-ferroptotic effects are likely to emerge.

### N-acetylcysteine protects cells from specific ferroptosis inducers and other electrophilic compounds

Organoid media formulations often include millimolar levels of N-acetylcysteine (NAC),^16^ a cell-permeable prodrug of cysteine. NAC is a poor lipophilic radical-trapping antioxidant^34^ but can be converted in cells into small molecules (e.g., glutathione) and proteins with antioxidant activity. The thiol moiety of NAC can also inactivate electrophilic small molecules via direct covalent reaction.^35^ We found that supplementing 2D culture medium with millimolar levels of NAC attenuated the potency of multiple lethal compounds, including both ferroptosis inducers and non-ferroptosis inducers (Figure 3A and Supplementary Figure 9). Among ferroptosis inducers, the potency of GPX4 inhibitors is likely attenuated because these compounds react with NAC at high concentration and prevent covalent inhibition of GPX4. In contrast, NAC may rescue from cyst(e)ine-depleting agents (e.g., erastin) by bypassing the requirement for system x_c_ ^−^ function and enabling direct uptake of NAC or mixed disulfides that are able to enter cells via alternate routes and are then converted intracellularly into cysteine.^36^

**Figure 3:**
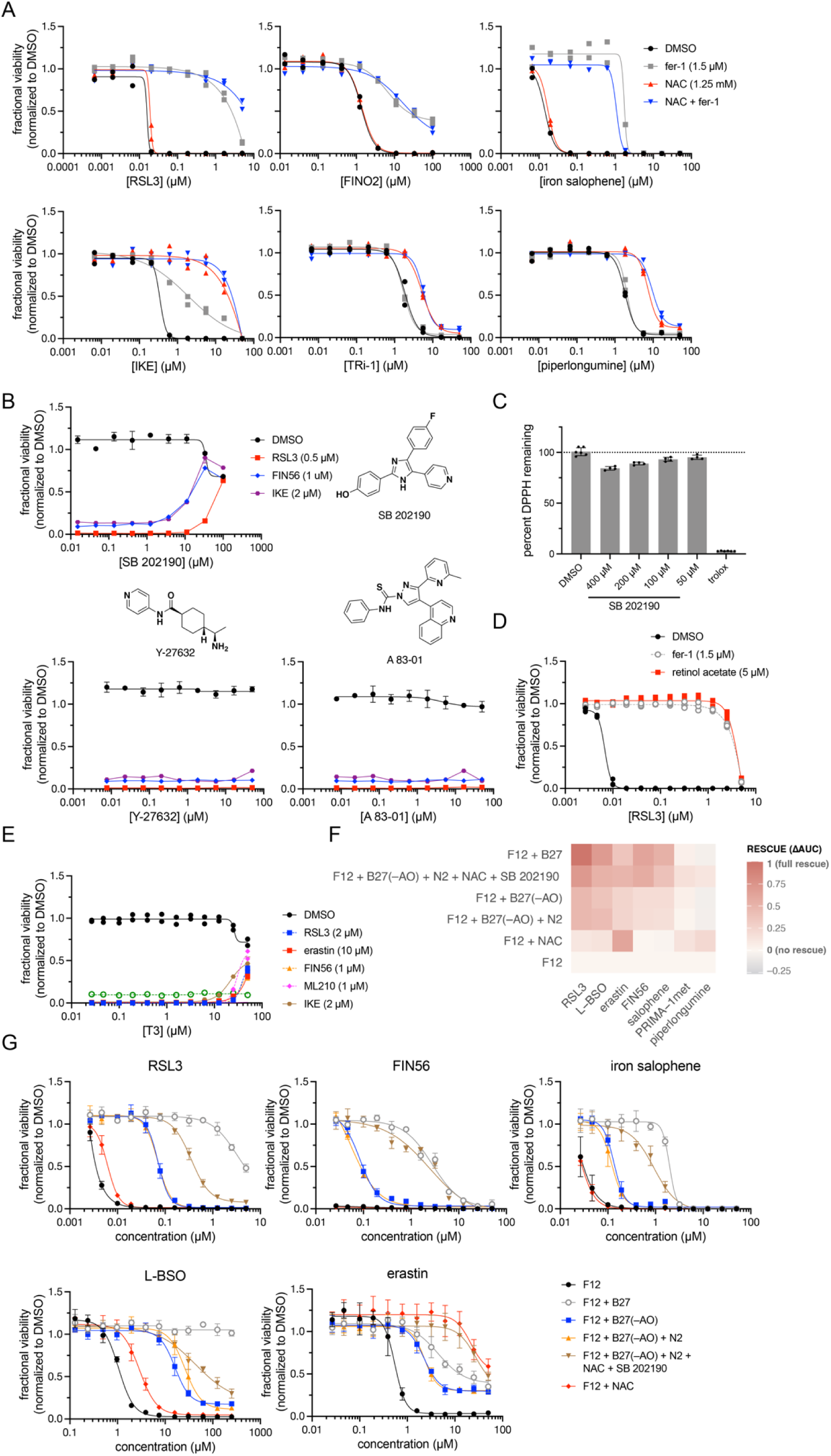
Small molecules used in organoid media protect cells from ferroptosis. (A) N-acetyl cysteine (NAC) protects cells from erastin and non-ferroptosis-inducing lethal compounds with electrophilic reactive groups. RKN cells were pretreated with ferrostatin (1.5 µM) and/or NAC (1.25 mM) for 24 h prior to treatment with the indicated compounds for 72 h. Data are plotted as two individual replicates. See also Supplementary Figure 9. (B) SB 202190 protects RKN cells from FINs at concentrations above 10 µM. Also shown are chemical structures and dose-response plots for SB 202190 and other small molecules commonly used in organoid medium formulations. Cells were treated with these small molecules followed by the treatment with the indicated ferroptosis inducers for 24 h prior to viability assessment. Data are plotted as mean ± s.d. for n = 3 replicates. (C) SB 202190 exhibits weak radical-trapping antioxidant activity as assessed by the DPPH assay. Compounds were incubated at ambient temperature for 1 h prior to absorbance measurements. Data are plotted as mean ± s.d. for n ≥ 4 replicates. (D) Vitamin A prodrug retinol acetate (5 µM) protects RKN cells from RSL3 (72 h treatment). Data are plotted as two individual replicates. (E) Thyroid hormone T3 (> 10 µM) protects cells from ferroptosis inducers. RKN cells were pretreated for 24 h with T3 followed by incubation with test compounds for 72 h. Data are plotted as two individual technical replicates. (F and G) The ferroptosis protective effects of medium supplements and other organoid components are additive. RKN cells were treated with the indicated medium supplements for 24 h followed by treatment with test compounds for 72 h. In panel F, heatmap colors are based on the difference in area-under-the-curve (ΔAUC) metric for viability of compound-treated cells relative to the F12 medium condition. In panel G, the same data is represented as dose–response curves, with data plotted as mean ± s.d. for n = 3 replicates.

However, the ability of NAC to protect cells against ferroptosis inducers was not universal. The potency of compounds that do not covalently inactivate GPX4 or inhibit glutathione biosynthesis was not affected by NAC treatment (Figure 3A). These observations highlight the importance of probing multiple ferroptosis-inducing perturbations in experiments seeking to draw conclusions about ferroptosis cell circuitry. The activity of non-ferroptosis inducers with electrophilic moieties was also suppressed by NAC treatment, including piperlongumine^37^, a molecule with multiple reactive sites, and Prima-1met^38^, a prodrug that forms a reactive ɑ,β-unsaturated ketone *in situ*. It is likely that such reactive compounds are inactivated by NAC directly or by increased levels of cellular glutathione following NAC treatment.

### Small-molecule pathway inhibitors used for organoid culture attenuate ferroptosis sensitivity

In addition to the additives discussed above, a variety of small molecules are commonly used to help generate organoid models, often at relatively high concentrations (1-10 µM) where effects may not be solely attributable to the annotated targets of these compounds. For example, it has been found that many small molecules possess intrinsic radical-trapping antioxidant activity or the ability to chelate iron and can thereby protect against ferroptosis, especially when used at high concentrations.^39,40^ We tested three compounds commonly used in organoid culture for such effects: Y-27632 (ROCK inhibitor), A 83-01 (ALK inhibitor), and SB 202190 (p38 MAPK inhibitor) (Figure 6A).

Y-27632 and A 83-01 did not have a significant effect on the potency of any ferroptosis inducers or lethal compound tested, even when tested at concentrations well above those used in media formulations (Figure 3B). SB 202190, however, was able to suppress the effects of multiple ferroptosis inducers when used at the reported concentration of 10 µM (Figure 3B). SB 202190 contains a phenol moiety that may be redox active and exhibits weak activity in the presence of DPPH, a stable radical used to measure compound radical-trapping antioxidant activity (Figure 3C). Consistent with our findings, SB 202190 was previously observed to suppress erastin-mediated ferroptosis in acute myeloid leukemia cells.^41^

In addition to synthetic small molecules, other vitamins and hormones present in organoid media formulations may also impact ferroptosis sensitivity. For example, vitamin A has been shown to modulate ferroptosis sensitivity through multiple proposed mechanisms.^42,43^ Vitamin A is included in B27 supplement as the prodrug form retinol acetate (Table 1), which is capable of rescuing cells from ferroptosis inducers at low micromolar concentrations (Figure 3D). The concentration of vitamin A in B27-supplemented medium is approximately 10-fold below the concentration required for ferroptosis suppression in RKN cells and omitting vitamin A from B27 (commercially available variant, B27–vitA) did not affect its ability to rescue from ferroptosis (Figure 2A). However, vitamin A may be a more effective ferroptosis inhibitor in other cell lines, and culturing cells in the presence of retinol acetate for longer periods may lead to the accumulation of vitamin A metabolites that more effectively protect against ferroptosis. Thyroid hormone T3 is another component of B27 supplement that has been demonstrated to be a radical-trapping antioxidant capable of inhibiting ferroptosis.^39^ At the recommended dilution of B27 supplement, the concentration of T3 is several orders of magnitude below that required to prevent ferroptosis in the models we tested. At high concentrations (>10 µM), T3 protects cells from GPX4 inhibitors and glutathione biosynthesis inhibitors (Figure 3E).

While molecules such as T3, vitamin A, and SB 202190 are less potent than widely used small-molecule ferroptosis inhibitors (such as ferrostatin-1, liproxstatin-1,^44^ and UAMC-3203^45^) and may not individually be present at concentrations sufficient for ferroptosis prevention, caution should be exercised when using media containing these compounds because the effects of small-molecule antioxidants are additive.^3^ Indeed, combined supplementation of basal medium with “antioxidant-free” variants B27–AO, N-2, NAC, and SB 202190 is sufficient to protect cells against ferroptosis inducers to a degree comparable to vitamin E-containing B27 or other ferroptosis inhibitors (Figures 3F and 3G).

### The anti-ferroptotic effects of organoid medium is reversible under certain conditions

Our findings raise the question of how ferroptosis can be studied in organoid models if such models depend on antioxidants in the culture medium for growth and maintenance. A potential solution is media replacement before ferroptosis-related experiments begin. To address this, we sought to determine whether media washout can reverse the anti-ferroptotic effects of organoid medium components following an initial period of exposure. In HT-1080 cells grown in 2D format, we found that the effects of B27 supplement on ferroptosis sensitivity persisted following removal of the medium, washing of cells with buffered saline, and addition of B27-free growth medium (Figure 4A). The magnitude of the residual protective effect of B27-treated cells increased with longer initial B27 exposure times. However, in other cases, the protective effects of commercial organoid medium could be eliminated through stringent washing (Figure 4B). It is unclear how extensible and effective this strategy of medium replacement can be for 3D models, where aggressive washing may be more difficult to achieve, and warrants further investigation.

**Figure 4:**
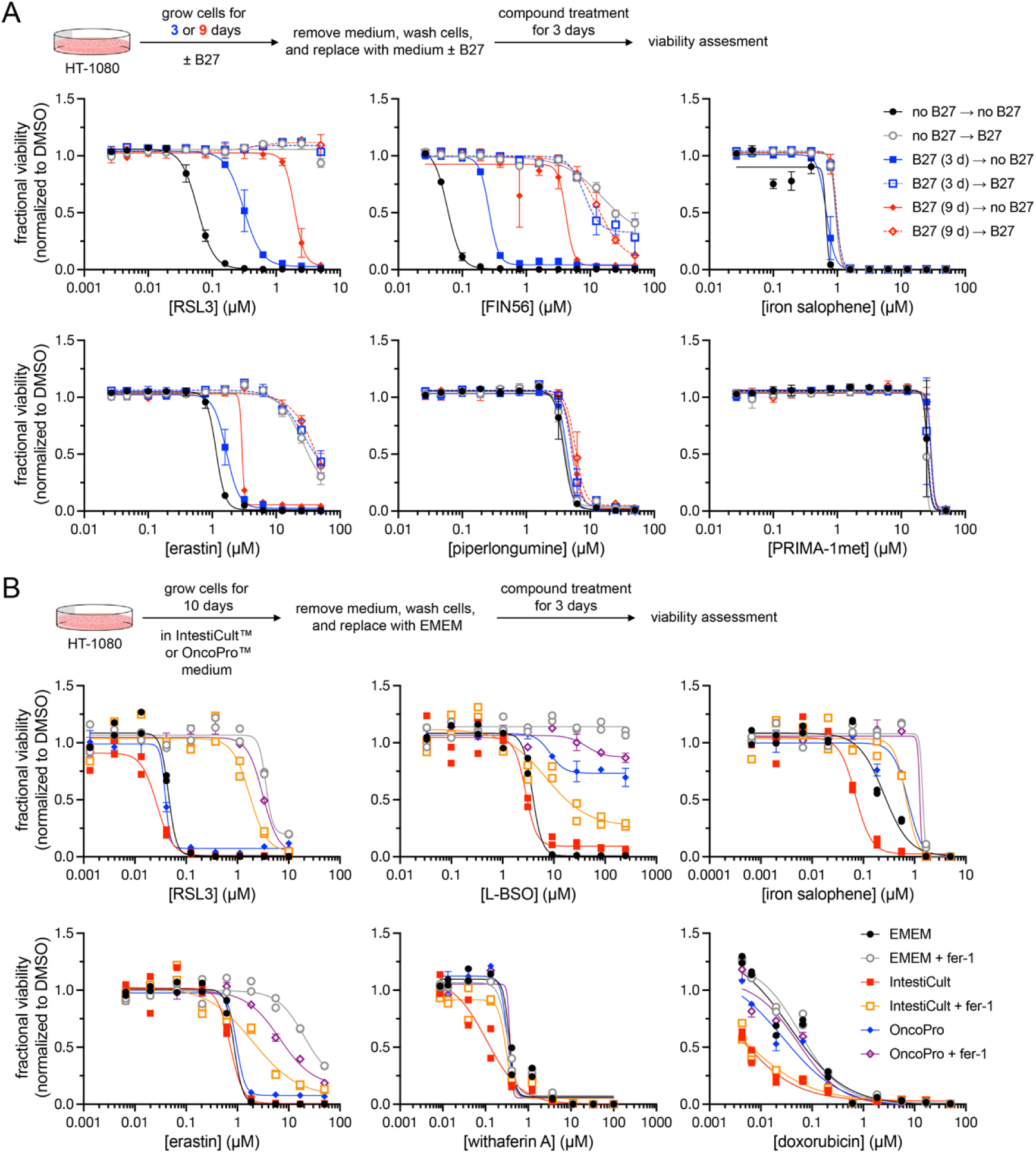
The anti-ferroptotic effects of organoid medium are reversible under certain conditions. (A) The ferroptosis-protective effect of B27 supplement in HT-1080 cells cannot be fully reversed by medium washout. Cells were cultured in EMEM with or without B27 (1x) for 3 or 9 days followed by medium removal, washing of cells with DPBS, and replacement of medium with or without B27 supplement. Cells were then treated with test compounds for 72 h prior to assessment of viability. Data are plotted as mean ± s.d. for n = 3 replicates. (B) The effects of the commercial IntestiCult™ and OncoPro™ media can be reversed in HT-1080 cells grown in 2D format. Cells were cultured in IntestiCult™ or OncoPro™ for 10 days, detached from the culture flask, washed with DPBS, and then plated in EMEM medium. Test compound treatment was performed for 72 h followed by viability assessment. Data are plotted as two individual technical replicates.

## Discussion

Uncontrolled lipid peroxidation is the central process that propels ferroptosis. The formation of lipid hydroperoxides depends on multiple factors including the availability of reactants (polyunsaturated lipids, iron, and oxygen), the basal level of cellular ROS generation, and cellular antioxidant capacity (protein and small-molecule antioxidants). Environmental factors can influence the availability of these substrates and modulate the balance between pro- and anti-oxidative forces within cells. Careful consideration of experimental design is required to assess ferroptosis given the complexity of processes that control cellular susceptibility to this form of death.

Our findings indicate many organoid media formulations, especially those utilizing B27 supplement, contain several lipophilic antioxidants, including but not limited to vitamin E, that suppress ferroptosis in both 2D and 3D culture. It is yet possible that other factors beyond lipophilic antioxidants may also impact cellular ferroptosis sensitivity. For example, in this study, we have neither examined the effects that protein and peptide growth factors used in organoid media have on ferroptosis sensitivity nor conditions that may alter transcriptional cell state and thereby ferroptosis sensitivity. Given the importance of employing organoids effectively for investigation of ferroptosis, we offer the following series of recommendations to select and validate media and organoid culture conditions that enable ferroptosis vulnerability to be probed:

- Test organoid medium in a ferroptosis-sensitive 2D cell model to determine the functional effect on ferroptosis; assess cell viability and ferroptosis biomarkers (e.g., lipid peroxidation).
- Identify culture conditions that may contain ferroptosis modulating components and exclude these from experimental workflows, if possible.
- If antioxidants are required for initial organoid model generation, determine if washout can be performed with a medium that does not inhibit ferroptosis before perturbational experiments are tested.
- Use multiple ferroptosis-inducing perturbations to assess ferroptosis sensitivity. For small molecules, test in dose at an appropriate concentration range and use a panel of structurally and mechanistically distinct inducers. Perform ferroptosis induction experiments with appropriate ferroptosis inhibitor controls (e.g., fer-1 and lip-1).
- Consider inclusion of multiple timepoints, as ferroptosis-inducing perturbations often exhibit characteristic kinetic profiles that differentiate them from other lethal agents.^40^

## Author contributions

J.K.E. and V.S.V. conceived of the project; J.K.E. designed and performed experiments; P.C., Y.Y.S., and R.A.V. supported cell culture maintenance and small molecule sensitivity profiling; P.C. performed 3D culture and *ex vivo* cell culture experiments; Y.Y.S. performed flow cytometry analyses; M.J.W. analyzed data and performed computational analyses; J.K.E., M.J.W. and V.S.V. wrote the paper.

---

**Supplementary Table 1.** Composition of B27 supplement and concentration of components at recommended dilution of 50x stock. Concentrations are estimated from Brewer and Cotman, *Brain Res.* 1989, 494, 65-74. The components in bold are antioxidants absent in B27–AO.

|  | component | 1x concentration (μM) |
| --- | --- | --- |
| 1 | BSA fraction V IgG free fatty acid poor | 37 |
| 2 | **catalase** | 0.001 |
| 3 | **glutathione (reduced)** | 3.2 |
| 4 | insulin (human) | 0.6 |
| 5 | **superoxide Dismutase** | 0.077 |
| 6 | holo-transferrin (human) | 0.062 |
| 7 | thyroid hormone T3 | 0.0026 |
| 8 | L-carnitine hydrochloride | 12 |
| 9 | ethanolamine | 16 |
| 10 | D-(+)-galactose | 83 |
| 11 | putrescine dihydrochloride | 183 |
| 12 | sodium selenite | 0.083 |
| 13 | corticosterone | 0.058 |
| 14 | linoleic acid | 3.5 |
| 15 | linolenic acid | 3.5 |
| 16 | progesterone | 0.02 |
| 17 | retinol acetate | 0.2 |
| 18 | **DL-alpha-tocopherol** | 2.3 |
| 19 | **DL-alpha-tocopherol acetate** | 2.1 |
| 20 | biotin | 0.4 |

**Supplementary Table 2.** Composition of N-2 supplement and concentration of components for Gibco™ formulation (ThermoFisher; catalog number: 17502048).

| Component | 100x concentration (μM) | 1x concentration (μM) |
| --- | --- | --- |
| holo-transferrin | 127 | 1.3 |
| insulin | 86 | 0.86 |
| progesterone | 2 | 0.02 |
| putrescine | 10000 | 100 |
| sodium selenite | 3 | 0.03 |

**Supplementary Figure 1:**
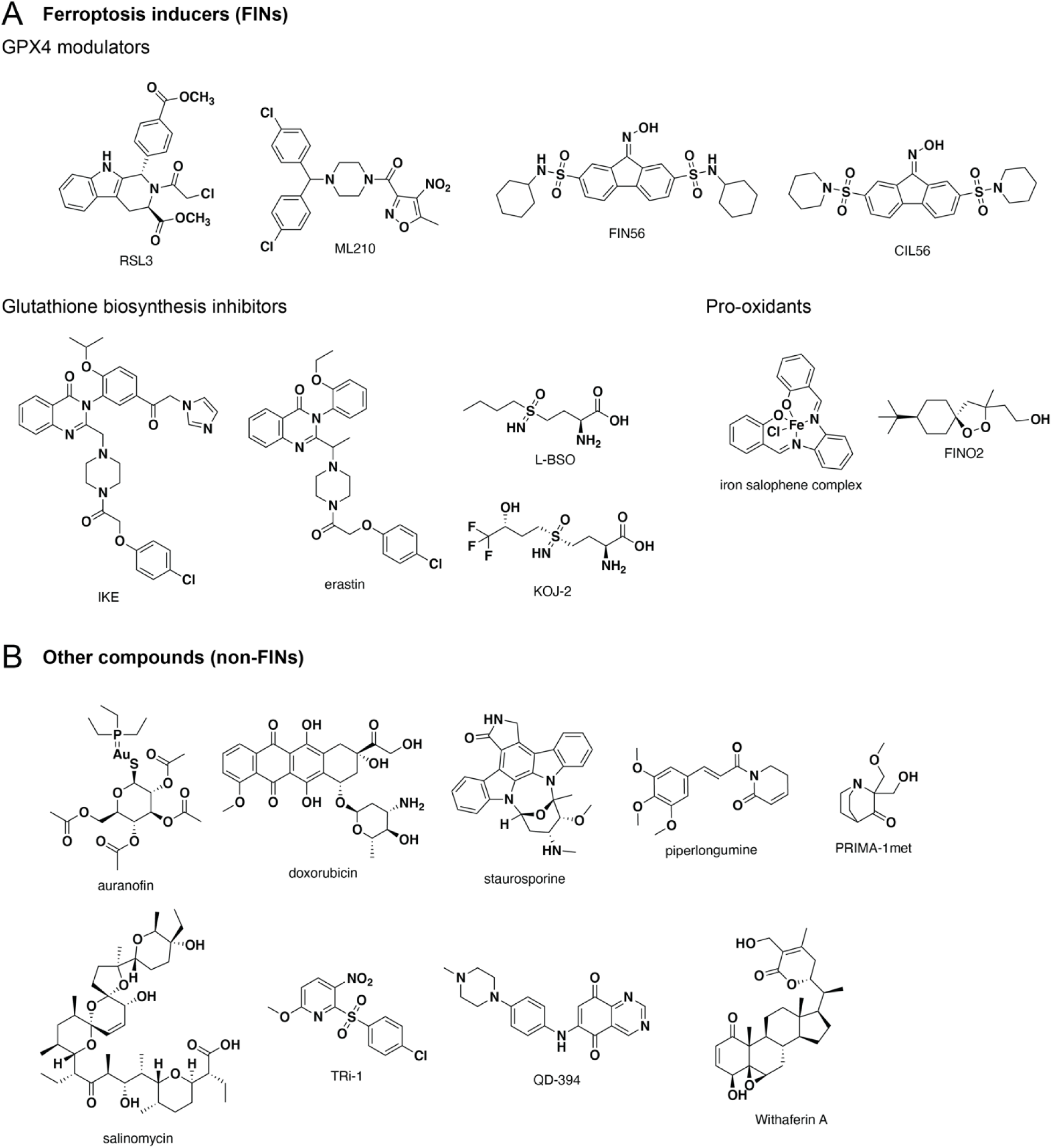
Chemical structures of compounds used in this study.

**Supplementary Figure 2:**
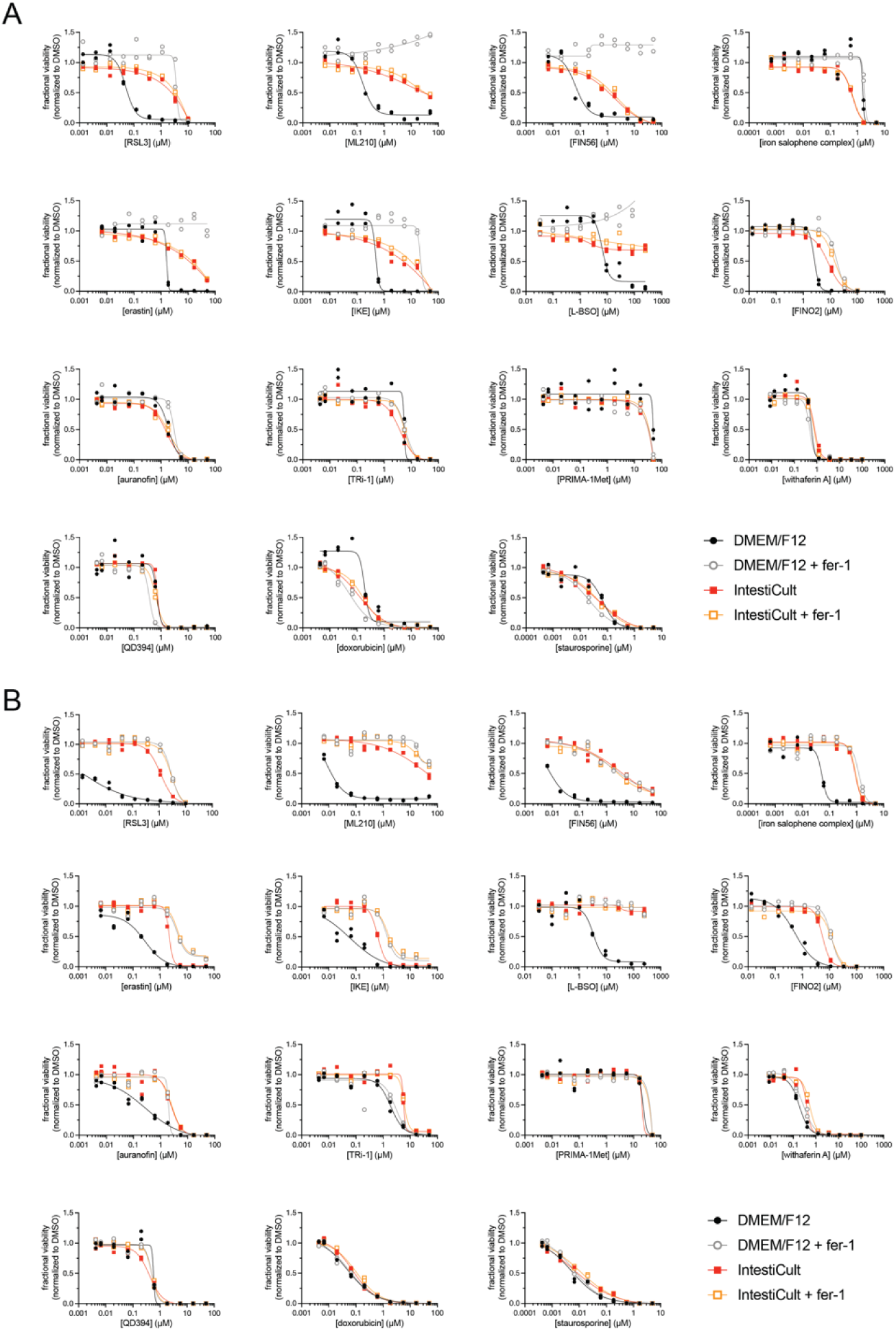
Entire dose-response curves for heatmaps in Fib 1B and 1D. IntestiCult™ medium protects cells from ferroptosis inducers in 2D (panel A) and 3D (panel B) culture formats. Compounds were incubated with cells for 72 h prior to viability assessment. Data are plotted as two individual technical replicates. Plots for RSL3 are reproduced from Figure 1.

**Supplementary Figure 3:**
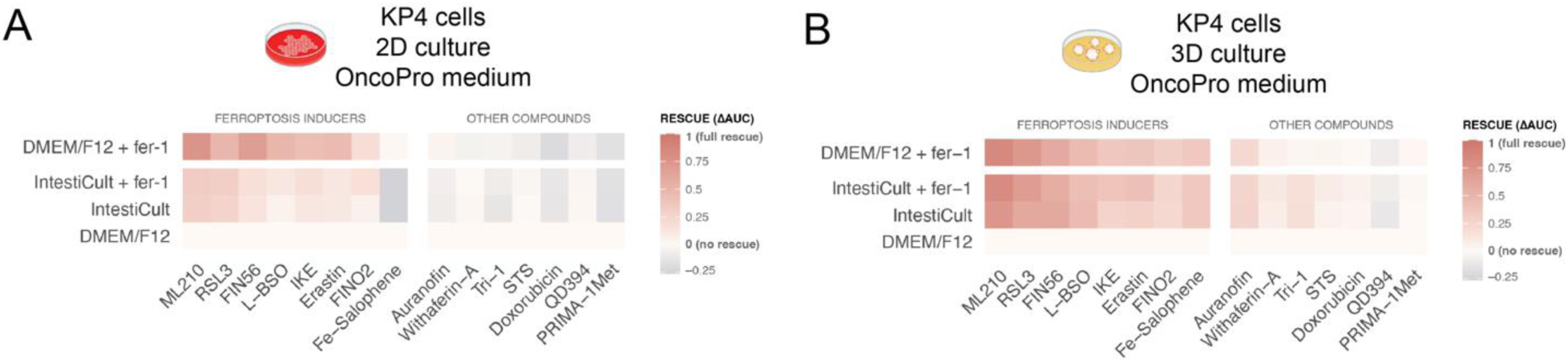
OncoPro medium suppresses ferroptosis in KP4 cells. OncoPro™ medium prevents ferroptosis in KP4 cells grown in 2D (panel A) and 3D (panel B) culture formats. Heatmap colors are based on the difference in area-under-the-curve (ΔAUC) metric for viability of compound-treated cells relative to the DMEM/F12 medium condition. AUC values for each compound relative to vehicle-treatment are derived from a 12-point dose response curve, n = 2 technical replicates. Compound treatment was performed for 72 h.

**Supplementary Figure 4:**
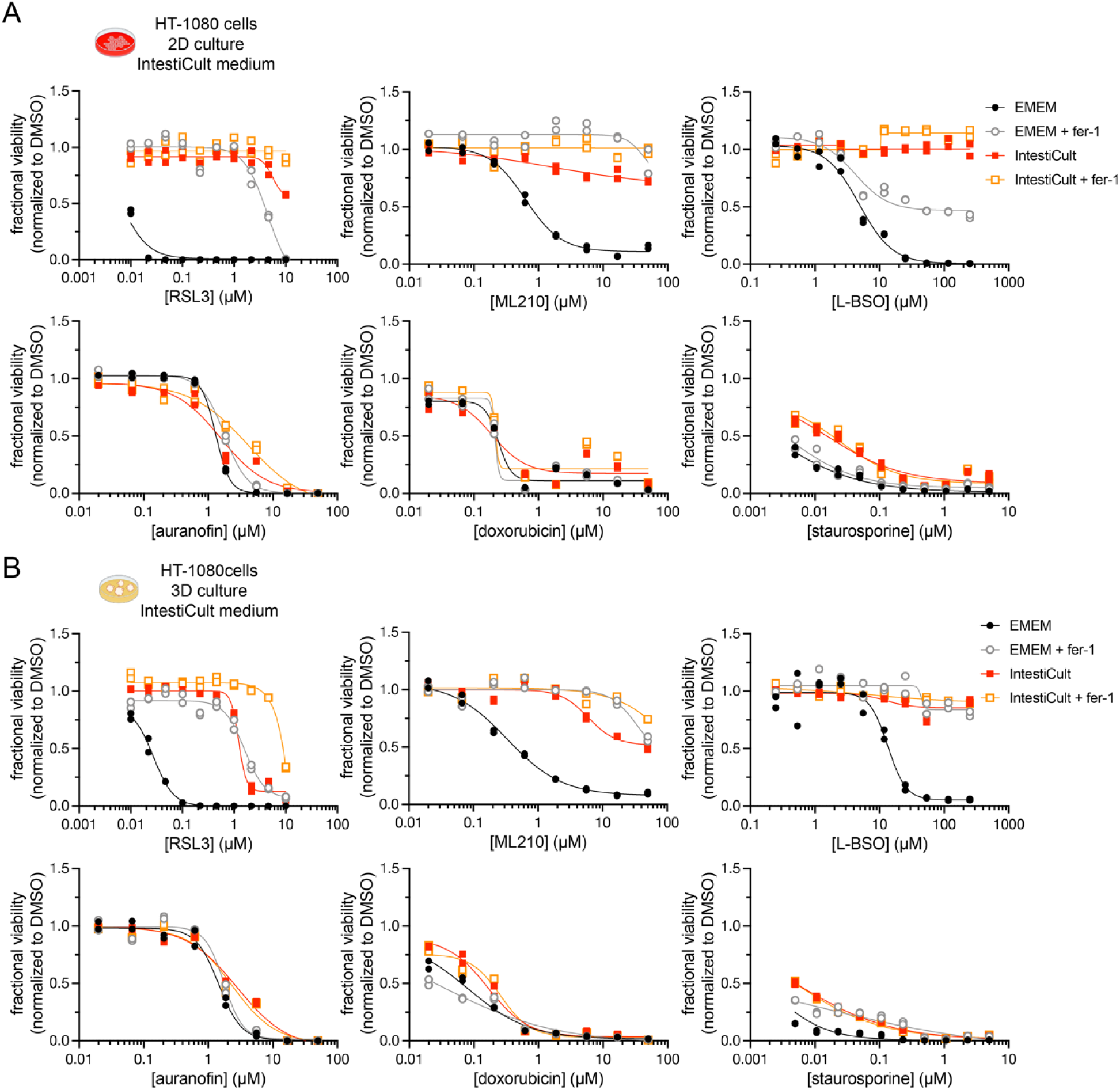
IntestiCult medium suppresses ferroptosis in HT-1080 cells. Ferroptosis could be prevented in HT-1080 cells by culturing in IntestiCult™ medium in 2D (panel A) and 3D (panel B) culture formats. Compounds were incubated with cells for 72 h prior to viability assessment. Data are plotted as two individual technical replicates.

**Supplementary Figure 5:**
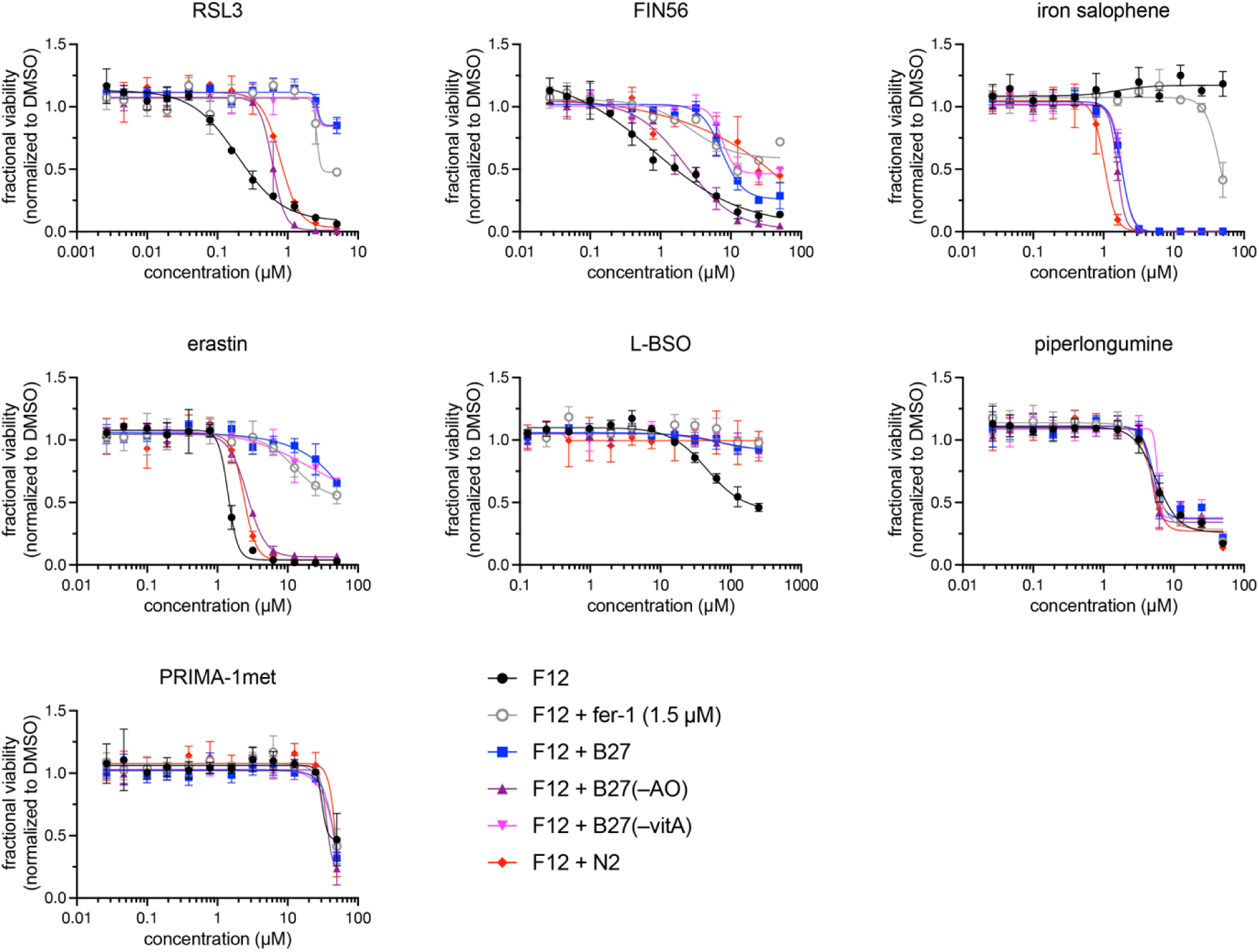
B27 supplement protects HT-1080 cells from ferroptosis inducers. B27 supplement protects HT-1080 cells from treatment with ferroptosis inducers (48 h). Data are plotted as mean ± s.d. for n = 3 replicates.

**Supplementary Figure 6:**
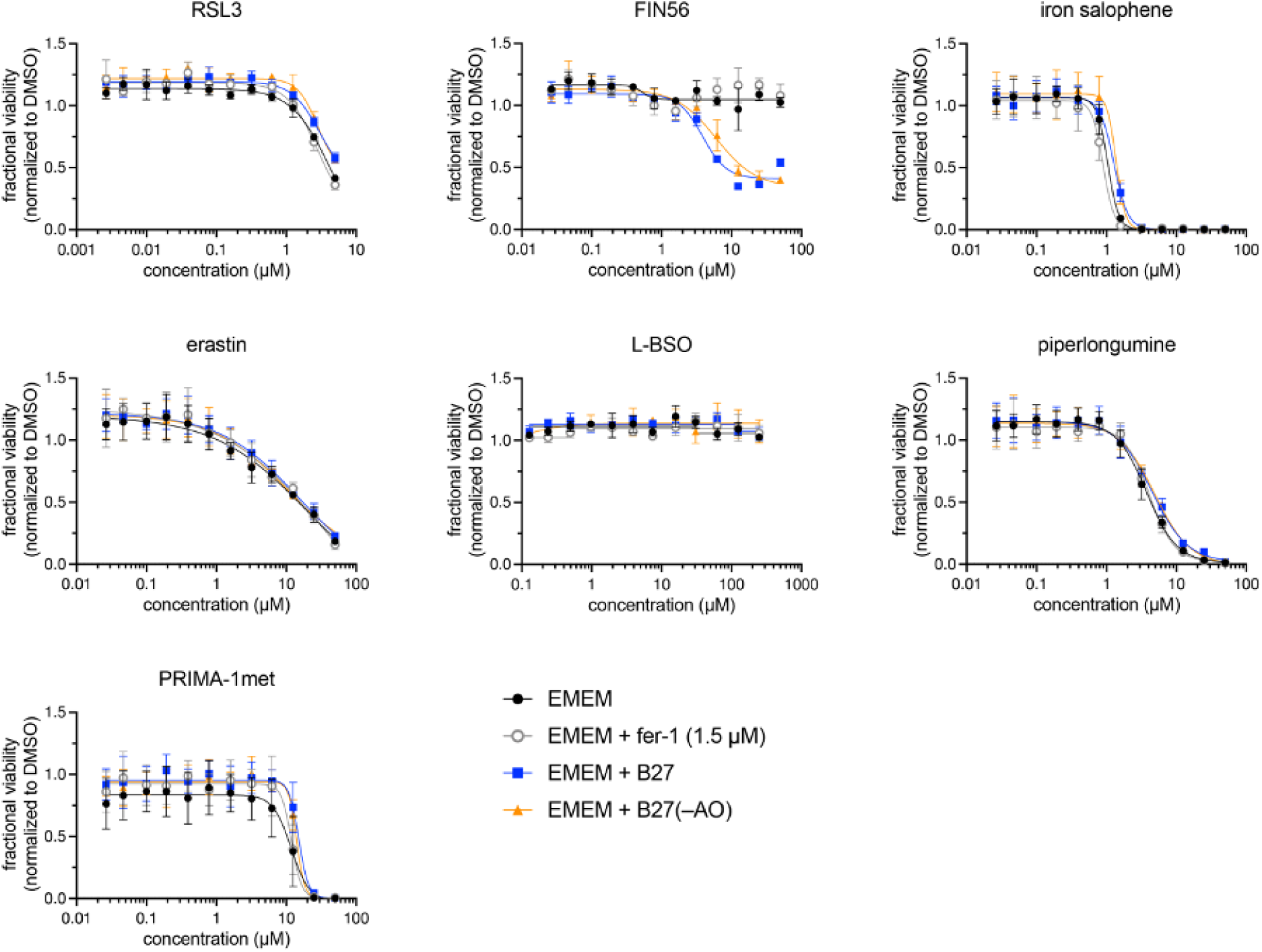
B27 supplement does not affect the sensitivity of a ferroptosis-insensitive cell line to ferroptosis inducers. B27 and B27(–AO) supplements do not protect MCF7 cells following treatment with lethal compounds for 48 h. Data are plotted as mean ± s.d. for n = 3 replicates.

**Supplementary Figure 7:**
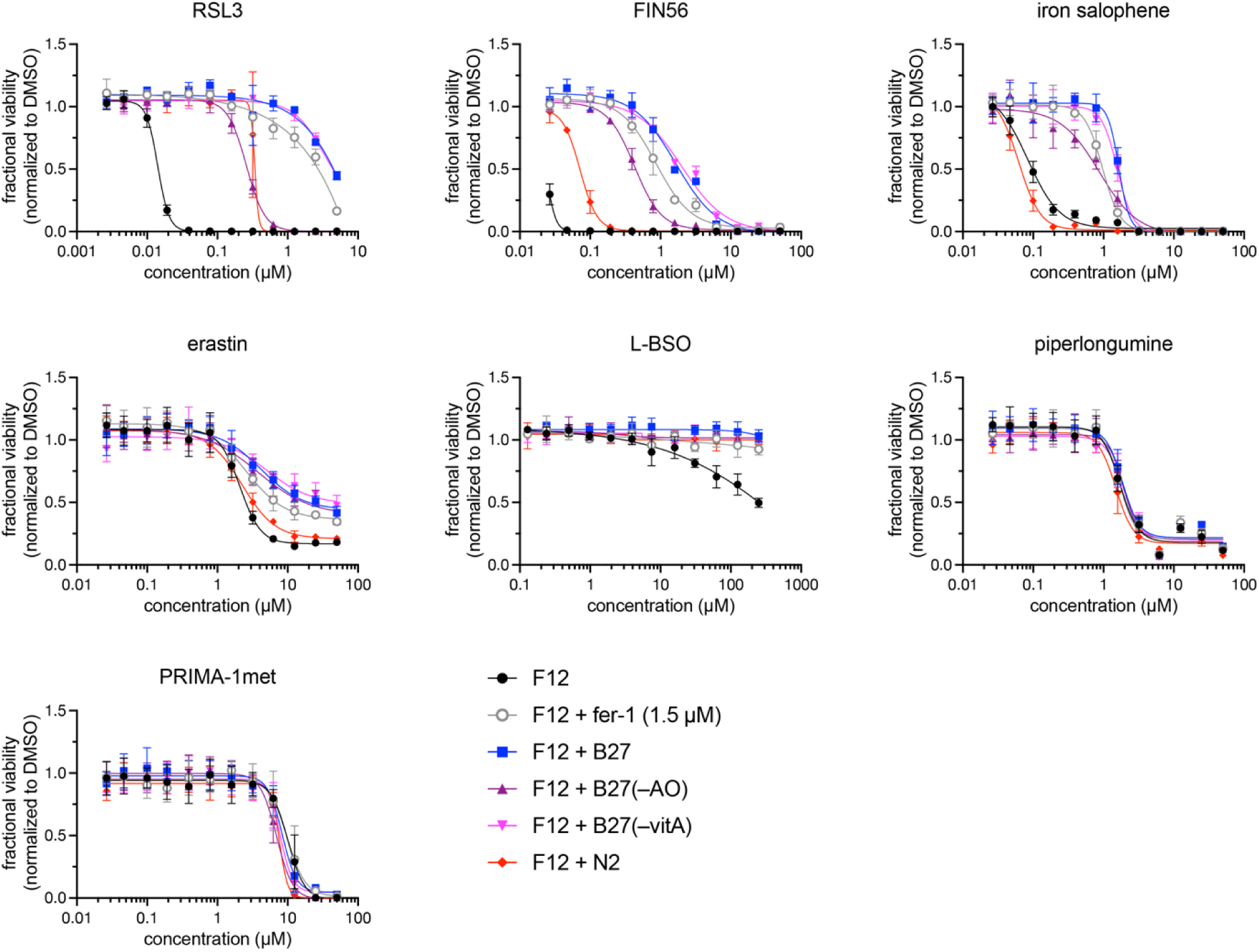
Entire dose-response curves for heatmap from Figure 2A. RKN cells cultured with the indicated media conditions were incubated with compounds for 48 h prior to viability assessment. Data are plotted as mean ± s.d. for n = 3 replicates. Related to Figure 2A.

**Supplementary Figure 8:**
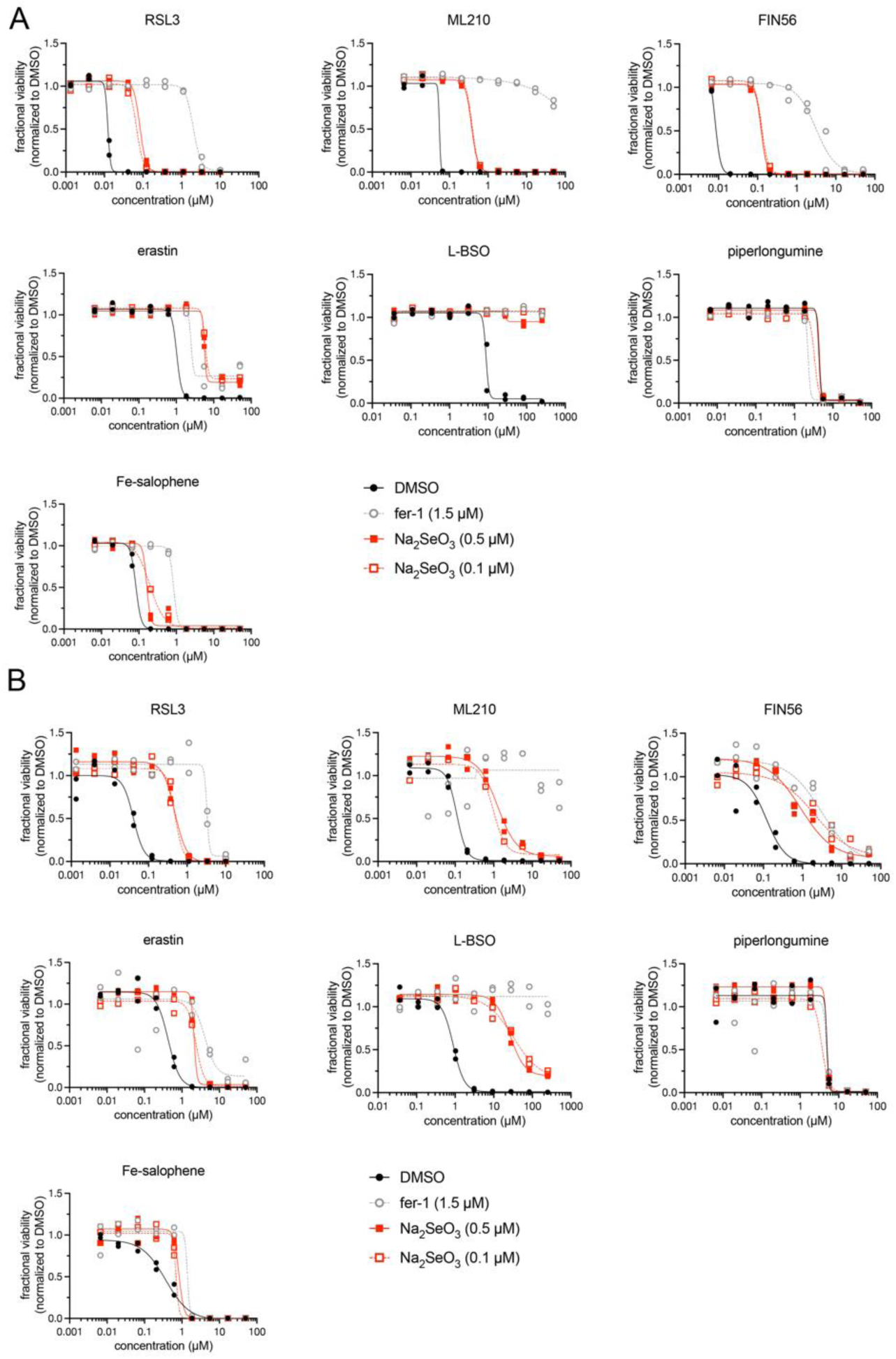
Sodium selenite protects RKN and HT-1080 cells from ferroptosis inducers. Pre-treatment of RKN (panel A) and HT-1080 (panel B) cells with sodium selenite for 24 h protected cells from ferroptosis inducers. RKN and HT-1080 cells were treated with compounds for 96 h prior to viability assessment. Data are plotted as two individual technical replicates. Related to Figure 2I.

**Supplementary Figure 9:**
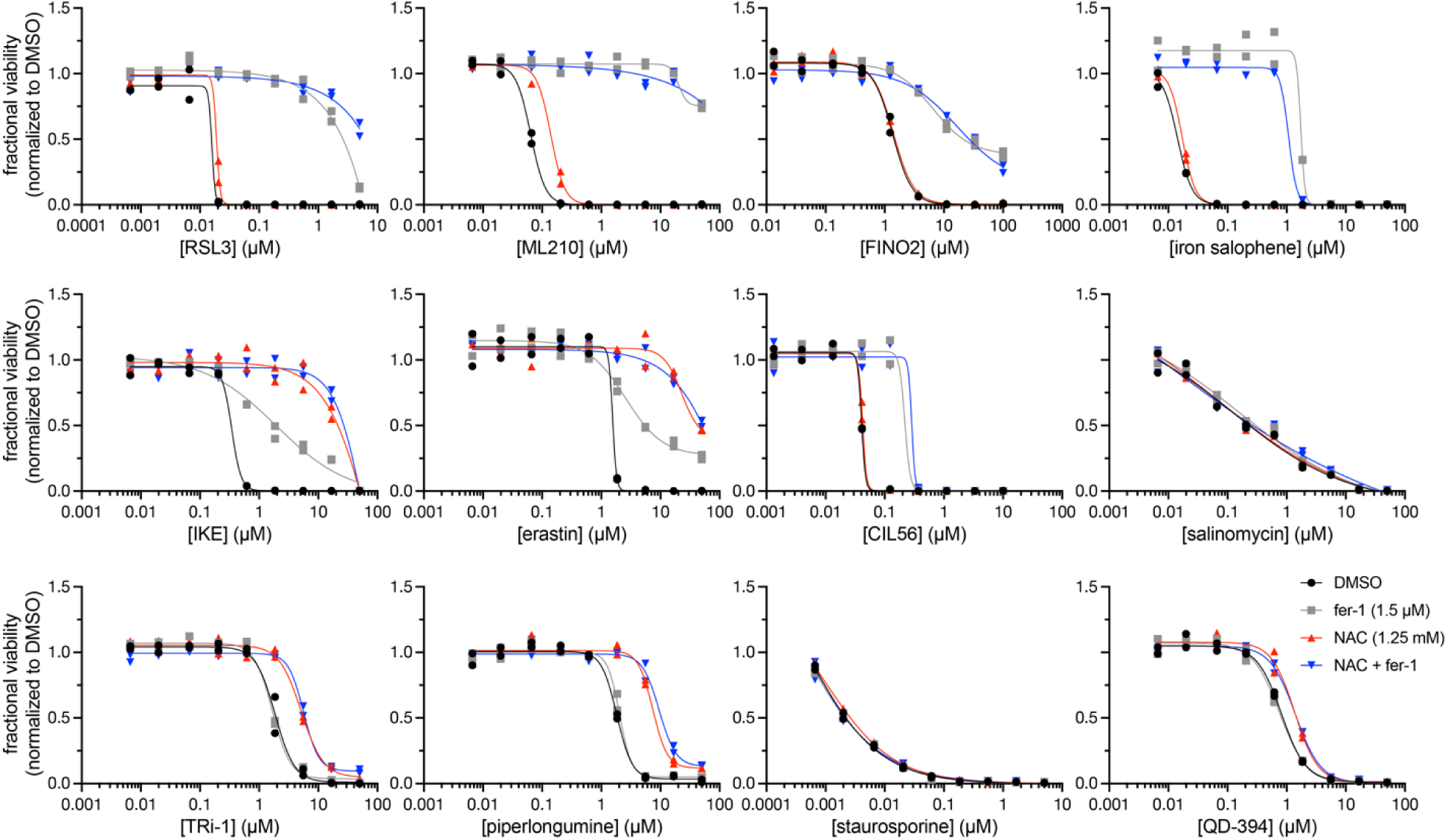
N-acetylcysteine protects cells from SLC7A11 inhibitors and non-FIN lethal compounds with reactive moieties. RKN cells were pretreated with ferrostatin (1.5 µM) and/or N-acetylcysteine (NAC; 1.25 mM) for 24 h prior to treatment with the indicated compounds for 72 h. Data are plotted as two individual replicates. Related to Figure 3A.

## References

1. Dixon, S. J. et al. Ferroptosis: An Iron-Dependent Form of Nonapoptotic Cell Death. Cell 149, 1060–1072 (2012).

2. Jiang, X., Stockwell, B. R. & Conrad, M. Ferroptosis: mechanisms, biology and role in disease. Nat. Rev. Mol. Cell Biol. 22, 266–282 (2021).

3. Dixon, S. J. & Pratt, D. A. Ferroptosis: A flexible constellation of related biochemical mechanisms. Mol. Cell 83, 1030–1042 (2023).

4. Yang, W. S., et al. Peroxidation of polyunsaturated fatty acids by lipoxygenases drives ferroptosis. Proc. Natl. Acad. Sci. 113, E4966–E4975 (2016).

5. Kagan, V. E. et al. Oxidized arachidonic and adrenic PEs navigate cells to ferroptosis. Nat. Chem. Biol. 13, 81–90 (2017).

6. Aldrovandi, M., Fedorova, M. & Conrad, M. Juggling with lipids, a game of Russian roulette. Trends Endocrinol. Metab. TEM 32, 463–473 (2021).

7. Yang, W. S. et al. Regulation of Ferroptotic Cancer Cell Death by GPX4. Cell 156, 317–331 (2014).

8. Doll, S. et al. FSP1 is a glutathione-independent ferroptosis suppressor. Nature 575, 693–698 (2019).

9. Bersuker, K. et al. The CoQ oxidoreductase FSP1 acts parallel to GPX4 to inhibit ferroptosis. Nature 575, 688–692 (2019).

10. Kraft, V. A. N. et al. GTP Cyclohydrolase 1/Tetrahydrobiopterin Counteract Ferroptosis through Lipid Remodeling. ACS Cent. Sci. 6, 41–53 (2020).

11. Ubellacker, J. M. et al. Lymph protects metastasizing melanoma cells from ferroptosis. Nature 585, 113–118 (2020).

12. Mishima, E. et al. A non-canonical vitamin K cycle is a potent ferroptosis suppressor. Nature 608, 778–783 (2022).

13. Barayeu, U. et al. Hydropersulfides inhibit lipid peroxidation and ferroptosis by scavenging radicals. Nat. Chem. Biol. 19, 28–37 (2023).

14. Freitas, F. P. et al. 7-Dehydrocholesterol is an endogenous suppressor of ferroptosis. Nature 626, 401–410 (2024).

15. Li, Y. et al. 7-Dehydrocholesterol dictates ferroptosis sensitivity. Nature 626, 411–418 (2024).

16. Urbischek, M. et al. Organoid culture media formulated with growth factors of defined cellular activity. Sci. Rep. 9, 6193 (2019).

17. Kim, J., Koo, B.-K. & Knoblich, J. A. Human organoids: model systems for human biology and medicine. Nat. Rev. Mol. Cell Biol. 21, 571–584 (2020).

18. Simian, M. & Bissell, M. J. Organoids: A historical perspective of thinking in three dimensions. J. Cell Biol. 216, 31–40 (2017).

19. Mayhew, C. N. & Singhania, R. A review of protocols for brain organoids and applications for disease modeling. STAR Protoc. 4, 101860 (2023).

20. Seiler, A. et al. Glutathione Peroxidase 4 Senses and Translates Oxidative Stress into 12/15-Lipoxygenase Dependent- and AIF-Mediated Cell Death. Cell Metab. 8, 237–248 (2008).

21. Skouta, R. et al. Ferrostatins Inhibit Oxidative Lipid Damage and Cell Death in Diverse Disease Models. J. Am. Chem. Soc. 136, 4551–4556 (2014).

22. Zilka, O. et al. On the Mechanism of Cytoprotection by Ferrostatin-1 and Liproxstatin-1 and the Role of Lipid Peroxidation in Ferroptotic Cell Death. ACS Cent. Sci. (2017).

23. Wu, J. et al. Intercellular interaction dictates cancer cell ferroptosis via NF2–YAP signalling. Nature 572, 402–406 (2019).

24. Bottenstein, J. E. & Sato, G. H. Growth of a rat neuroblastoma cell line in serum-free supplemented medium. Proc. Natl. Acad. Sci. 76, 514–517 (1979).

25. Brewer, G. J. & Cotman, C. W. Survival and growth of hippocampal neurons in defined medium at low density: advantages of a sandwich culture technique or low oxygen. Brain Res. 494, 65–74 (1989).

26. Brewer, G. J., Torricelli, J. R., Evege, E. K. & Price, P. J. Optimized survival of hippocampal neurons in B27-supplemented neurobasal^TM^, a new serum-free medium combination. J. Neurosci. Res. 35, 567–576 (1993).

27. Southon, A. et al. Cu ^II^ (atsm) inhibits ferroptosis: Implications for treatment of neurodegenerative disease. Br. J. Pharmacol. 177, 656–667 (2020).

28. Else, P. L. The highly unnatural fatty acid profile of cells in culture. Prog. Lipid Res. 77, 101017 (2020).

29. Eaton, J. K., et al. The enzyme glutamate-cysteine ligase (GCL) is a target for ferroptosis induction in cancer. bioRxiv (2024).

30. Shimada, K. et al. Global survey of cell death mechanisms reveals metabolic regulation of ferroptosis. Nat. Chem. Biol. 12, 497–503 (2016).

31. Tuo, Q.-Z. et al. Characterization of Selenium Compounds for Anti-ferroptotic Activity in Neuronal Cells and After Cerebral Ischemia–Reperfusion Injury. Neurotherapeutics 18, 2682– 2691 (2021).

32. Li, Z., et al. Ribosome Stalling during Selenoprotein Translation Exposes a Ferroptosis Vulnerability in Cancer. http://biorxiv.org/lookup/doi/10.1101/2022.04.11.487892(2022) doi:10.1101/2022.04.11.487892.

33. Lee, N. et al. Selenium reduction of ubiquinone via SQOR suppresses ferroptosis. Nat. Metab. 6, 343–358 (2024).

34. Shah, R., Farmer, L. A., Zilka, O., Van Kessel, A. T. M. & Pratt, D. A. Beyond DPPH: Use of Fluorescence-Enabled Inhibited Autoxidation to Predict Oxidative Cell Death Rescue. Cell Chem. Biol. 26, 1594–1607.e7 (2019).

35. Adams, D. J. et al. Discovery of Small-Molecule Enhancers of Reactive Oxygen Species That are Nontoxic or Cause Genotype-Selective Cell Death. ACS Chem. Biol. 8, 923–929 (2013).

36. Dixon, S. J. et al. Pharmacological inhibition of cystine–glutamate exchange induces endoplasmic reticulum stress and ferroptosis. eLife 3, e02523 (2014).

37. Adams, D. J. et al. Synthesis, cellular evaluation, and mechanism of action of piperlongumine analogs. Proc. Natl. Acad. Sci. 109, 15115–15120 (2012).

38. Lambert, J. M. R. et al. PRIMA-1 Reactivates Mutant p53 by Covalent Binding to the Core Domain. Cancer Cell 15, 376–388 (2009).

39. Mishima, E. et al. Drugs Repurposed as Antiferroptosis Agents Suppress Organ Damage, Including AKI, by Functioning as Lipid Peroxyl Radical Scavengers. J. Am. Soc. Nephrol. 31, 280–296 (2020).

40. Conlon, M. et al. A compendium of kinetic modulatory profiles identifies ferroptosis regulators. Nat. Chem. Biol. 17, 665–674 (2021).

41. Yu, Y. et al. The ferroptosis inducer erastin enhances sensitivity of acute myeloid leukemia cells to chemotherapeutic agents. Mol. Cell. Oncol. 2, e1054549 (2015).

42. Tschuck, J., et al. Suppression of ferroptosis by vitamin A or antioxidants is essential for neuronal development. bioRxiv (2023) doi:10.1101/2023.04.05.535746.

43. Jakaria, Md., Belaidi, A. A., Bush, A. I. & Ayton, S. Vitamin A metabolites inhibit ferroptosis. Biomed. Pharmacother. 164, 114930 (2023).

44. Friedmann Angeli, J. P., et al. Inactivation of the ferroptosis regulator Gpx4 triggers acute renal failure in mice. Nat. Cell Biol. 16, 1180–1191 (2014).

45. Devisscher, L. et al. Discovery of Novel, Drug-Like Ferroptosis Inhibitors with in Vivo Efficacy. J. Med. Chem. 61, 10126–10140 (2018).

